# A role for glial fibrillary acidic protein (GFAP)-expressing cells in the regulation of gonadotropin-releasing hormone (GnRH) but not arcuate kisspeptin neuron output

**DOI:** 10.1101/2021.03.10.434805

**Authors:** Charlotte Vanacker, R. Anthony DeFazio, Charlene M. Sykes, Suzanne M. Moenter

**Author notes:** Corresponding author, Suzanne M Moenter, Ph.D., Department of Molecular and Integrative Physiology, University of Michigan, 7725 Med Sci II, 1137 E. Catherine St, Ann Arbor, MI 48109-5622, (voice).

## Abstract

GnRH neurons are the final central neural output regulating fertility. Kisspeptin neurons in the hypothalamic arcuate nucleus (KNDy neurons) are considered the main regulator of GnRH output. GnRH and KNDy neurons are surrounded by astrocytes, which can modulate neuronal activity and communicate over distances. Prostaglandin E2 (PGE2), synthesized primarily by astrocytes, increases GnRH neuron activity and downstream pituitary release of luteinizing hormone (LH). We hypothesized GFAP-expressing astrocytes play a role regulating GnRH and/or KNDy neuron activity and LH release. We used adenoassociated viruses to target designer receptor exclusively activated by designer drugs (DREADDs) to GFAP-expressing cells to activate Gq or Gi-mediated signaling. Activating Gq signaling in the preoptic area, near GnRH neurons, but not in the arcuate, increases LH release *in vivo* and GnRH firing *in vitro* via a mechanism in part dependent upon PGE2. These data suggest astrocytes can activate GnRH/LH release in a manner independent of KNDy neurons.

## Introduction

Reproduction is controlled by the hypothalamo-pituitary-gonadal axis. The episodic release of gonadotropin-releasing hormone (GnRH) from neurons in the preoptic area (POA) of the hypothalamus drives the function of this axis.^1^ GnRH stimulates the anterior pituitary gland to synthesize and secrete two gonadotropins, LH and follicle-stimulating hormone (FSH), which subsequently activate gonadal functions including steroidogenesis. Sex steroids feed back to regulate the GnRH/LH release in both sexes ^2–5^.

How GnRH neurons are activated remains incompletely understood. Over the past several years, kisspeptin neurons located in the arcuate nucleus of the hypothalamus (ARC) have emerged as a leading candidate for initiating GnRH/LH release^6–10^, as well as for integrating steroid negative feedback ^11–14^. These cells are called KNDy neurons for their coexpression of kisspeptin, neurokinin B and dynorphin. KNDy neurons in brain slices exhibit changes in spontaneous action potential firing activity that is on the time-scale of episodic LH release and that is steroid sensitive^13^. *In vivo,* optogenetic activation of KNDy neurons induces LH release, and increases in intracellular calcium in these cells measured by bulk photometry are associated with spontaneous LH release^10,15^. Whether or not KNDy neurons endogenously generate the release of GnRH or are a relay station for a signal generated in other cell types is not known, nor is the list of other cell types that may alter GnRH neuron firing independent of KNDy neurons complete.

In this regard, evidence suggests astroglia play a role in reproductive function^16–21^. Astrocytes are a major glial type in the central nervous system. GnRH and KNDy neurons are contacted by astrocytes and astrocytic coverage of neurons in the POA and ARC varies with estrous cycle stage, gonadal status, and seasonal reproduction^22–26^. Astrocytes can propagate information to adjacent astrocytes^27–29^ making them intriguing candidates for coordinating episodic hormone release over the distances among scattered GnRH neurons. A consensus has yet to emerge regarding vesicle-mediated transmission by astrocytes^30^, but there are data indicating these cells may release multiple gliotransmitters^31–33^. Among these is PGE2, a non-vesicular transmitter that can increase GnRH neuron activity, GnRH release and LH release ^34–36^.

Our working hypothesis is that local astrocytes modulate GnRH and/or KNDy neuron activity to alter LH release *in vivo.* To investigate the role of astrocyte signaling in the control of GnRH and KNDy neurons *in vitro* and LH release *in vivo,* we took advantage of designer receptors exclusively activated by designer drugs (DREADDs). DREADDs were genetically targeted to astroglia by putting them under the control of the glial fibrillary acidic protein (GFAP) promoter coupled with local adeno-associated virus (AAV) delivery. GFAP is mainly expressed by astrocytes in the central nervous system^37^, thus this approaches allowed temporal and spatial control of G-protein-coupled receptor (GPCR) signaling in astrocytes. Our results indicate that activating signaling in GFAP-expressing cells can induce GnRH neuron activity and LH release in a sex-dependent manner, but does not alter KNDy neuron activity. This suggests central elements other than KNDy neurons can initiate GnRH release.

## Results

### AA V5 bearing GFAP promoter-driven construct effectively targets GFAP-expressing cells

To examine the response of the reproductive neuroendocrine system to activation of Gq signaling in GFAP-expressing cells we injected AAV5 bearing GFAP promoter-driven constructs expressing either the mCherry reporter or Gq- or Gi-coupled DREADDs and mCherry. Table 1 shows the animal models and viruses used in this work and how they are abbreviated in the text, Table 2 shows the primary antibodies). GFAP is expressed in glial cells throughout the body, including, for example, the pancreas and pituitary^38,39^. We thus used stereotaxic injection to limit manipulations to the region of interest. Bilateral viral injections were made in adult mice into either the POA near GnRH soma (Figure 1A,C), or ARC near KNDy neurons (Figure 1B,D). Successful infection of the area of interest was confirmed by localized expression of mCherry (Figure 1A, B). Dual immunofluorescence for mCherry and either the astroglial cytoplasmic marker S100β (Figure 1E), or the neuronal marker NeuN (Figure 1F), were performed to characterize the infected population. S100β was used to identify astroglia as GFAP was the driving promoter in the viral constructs used. S100β and GFAP are colocalized in most (<90%) cells examined hypothalamic tissue (Supplemental Figure 1). Data for colocalization of S100β and NeuN with mCherry were similar between ARC and POA and were combined for analysis. Over 90% of mCherry infected cells colocalized with S100β regardless of virus used or region examined (Figure 1G). A small percent of mCherry infected cells colocalized with NeuN and about 2% of mCherry infected cells did not colocalize with either marker. DREADD expression was thus primarily targeted to astrocytes and compatible with *in vitro* and *in vivo* approaches. Because of the small percent of neuronal colocalization, all slices for physiology studies were examined for mCherry infection of neurons either based on morphology or NeuN staining. No expression of mCherry was observed in any recorded or imaged GnRH-GFP or Tac2-GFP neuron, thus cells from which data were directly obtained were likely not infected. Data from mice (for LH measures) or from those brain slices with mCherry signal was observed in cells with neuronal morphology (for electrophysiology and Ca measurements) were excluded from statistical analyses. For transparency, these data are shown in different colors (magenta for unidentified neurons, cyan for Tac2-expressing neurons) when individual cell data are presented.

**Table 1.** Mouse strains and viruses used

|  | Full name | referred to in text as | reference | Supplier catalog # | RRID |
| --- | --- | --- | --- | --- | --- |
| mouse | B6;CBATg(Gnrh1-EGFP)51Sumo/J | GnRH-GFP | <sup>66</sup> | JAX 033639 | IMSR_JAX:033639 |
| mouse | Tg [Tac2-EGFP]381Gsat | Tac2-GFP | <sup>67</sup> | MMRRC; 015495-UCD/STOCK | MMRRC_015495-UCD |
| AAV | AAV5-GFAP-hM3Dq(Gq)-mCherry | AAV-Gq | <sup>73</sup> | Addgene; 50478-AAV5 | Addgene_50478 |
| AAV | AAV5-GFAP-hM4Di(Gi)-mCherry | AAV-Gi | This paper | Addgene;50479-AAV5 | Addgene_50479 |
| AAV | AAV5-GFAP104-mCherry | AAV-mCherry | This paper | Addgene; 58909-AAV5 | Addgene_58909 |
| AAV | AAV5-gfaABC1D-cyto-GCaMP6f | AAV-GCaMP6f | <sup>74</sup> | Addgene;52925-AAV5 | Addgene_52925 |

**Table 2.** Antibodies used

| Antibody | species | dilution/use | reference | Supplier catalog # | RRID |
| --- | --- | --- | --- | --- | --- |
| Anti-HA | rat | 1:3000 |  | Roche, clone 3F10 11867423001 | RRID:AB 390918 |
| Anti-GFAP | rabbit | 1:10000 |  | Agilent DAKO z0334 | RRID:AB 10013382 |
| Anti-S100β | mouse | 1:500 | | Sigma S2532 | RRID:AB_477499 |
| Anti-mCherry | rat | 1:3000 |  | Life Technologies M11217 | RRID:AB_2536611 |
| Anti-NeuN | rabbit | 1:3000 |  | Abcam ab177487 | RRID:AB 2532109 |
| Anti-GFP | chicken | 1:1000 |  | Abcam ab13970 | RRID:AB_300798 |
| anti-bovine LHβ 518B7 | | LH assay capture | | Janet Roser, UC Davis | RRID AB_2665514 |
| AFP240580Rb | rabbit | LH assay detection |  | National Hormone & Peptide Program | RRID AB_2665533 |
| goat anti-rabbit HRP | goat | LH assay enzyme conjugated |  | Dako D048701-2 |  |

**Figure 1.**
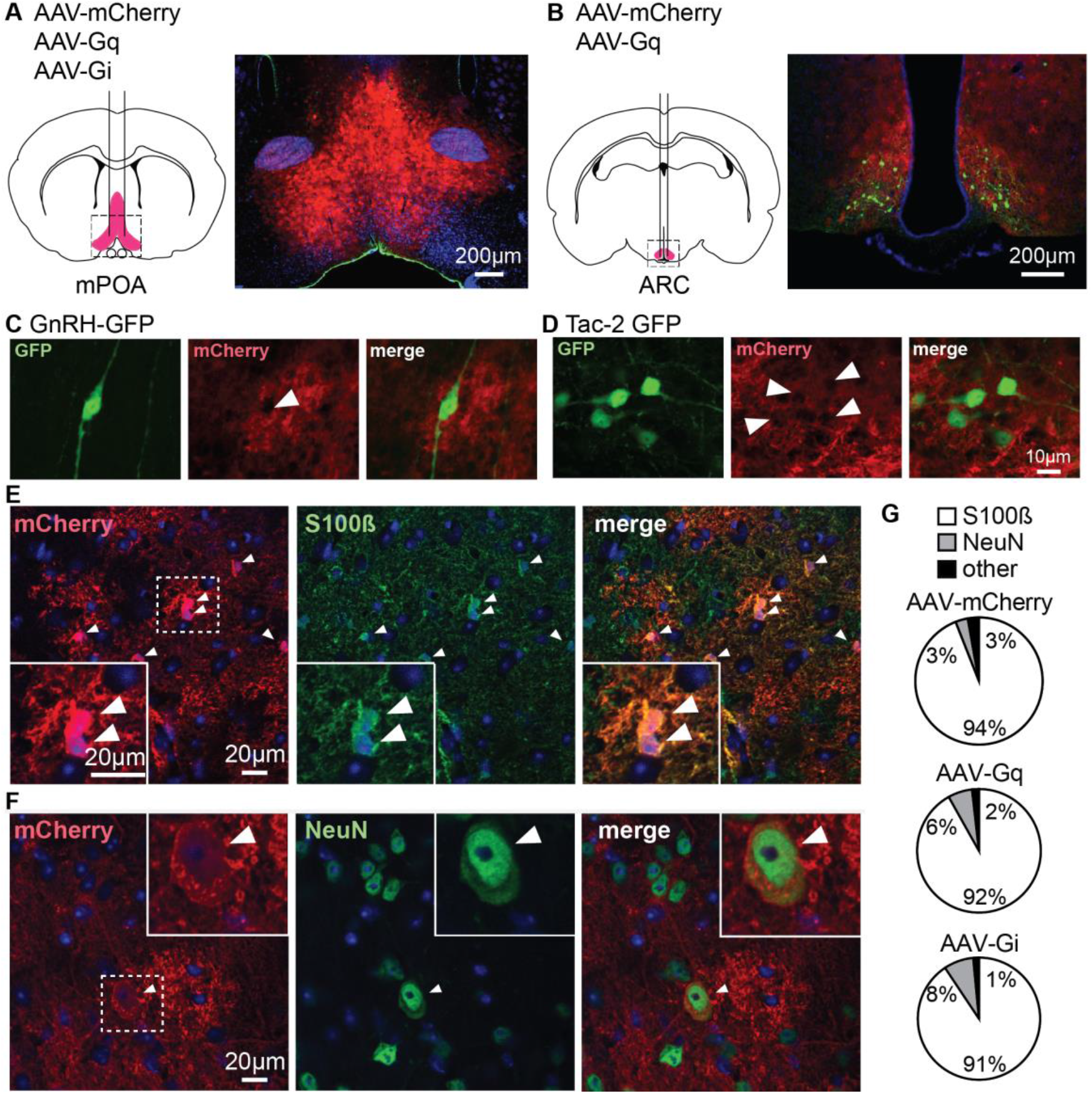
AAV5 bearing GFAP promoter-driven constructs effectively target primarily cells with astroglial phenotype and markers. A,B. Bilateral stereotaxic injection (left) and corresponding infection (right) of the POA (A) and ARC (B). C,D. Dual immunofluorescence for GFP (left) and mCherry (middle) in the POA (C) and ARC (D) reveals GFP-positive neurons surrounded by infected tissue (merge, right). Arrowheads in middle panels of C, D show gaps in mCherry signal where neurons are located. E. Immunofluorescence for mCherry (left), S100β (middle) and merge (right) showing colocalization of the two signals, white arrowheads identify colocalization between mCherry and S100β staining; dashed box shows area magnified in lower left. F. Immunofluorescence for mCherry (left), NeuN (middle) and merge (right) showing colocalization of the two signals, white arrowheads identify colocalization between mCherry and NeuN staining; dashed box shows area magnified in upper right. G. Quantification of infected cells expressing specific markers for each virus type.

### Activation of Gq but not Gi signaling in GFAP-expressing cells alters intracellular calcium levels

Activation of GPCRs including DREADDs increases intracellular calcium in astrocytes from different brain areas including the hypothalamus^40,41^. To characterize the effect of DREADD activation on intracellular calcium in our model, we injected wild-type male mice with an AAV5 encoding the calcium indicator GCaMP6f under the control of GFAP promoter (AAV-GCaMP6f) together with the either AAV-Gq or AAV-Gi or AAV-mCherry control virus (Table 3). Because LH release and GnRH neuron activity can be activated by PGE2, which is one putative gliotransmitter^42^, we postulated that activation of Gq signaling in GFAP-expressing cells would be activating to the reproductive neuroendocrine system, whereas activation of Gi would be inhibitory. Gonad-intact mice were thus used in tests of Gq activation, and a mix of gonad-intact and castrated mice (to increase activity of the reproductive neuroendocrine system) in tests of Gi activation.

**Table 3.** Co-injected viruses, regions, gonadal status for calcium imaging studies of male mice co-infected with AAV-GCaMP6f. P <0.05 is shown in bold.

| virus | region | gonadal status | # mice | # recordings | ANOVA | F | P |
| --- | --- | --- | --- | --- | --- | --- | --- |
| AAV-mCherry | POA | intact | 1 | 3 | virus | (2,8)=5.616 | **0.0299** |
| AAV-Gq | POA | intact | 2 | 5 | treatment | (1.432,11.46)=11.68 | **0.0031** |
| AAV-Gi | POA | intact | 1 | 3 | interaction | (4,16)=11.49 | **0.0001** |
| AAV-mCherry | POA | castrated | 1 | 3 | virus | (1,7)=0.3861 | 0.5540 |
| AAV-Gi | POA | castrated | 2 | 6 | treatment | (2,14)=0.4036 | 0.6754 |
|  |  |  |  |  | interaction | (2,14)=1.417 | 0.2751 |
| AAV-mCherry | ARC | intact | 1 | 3 | virus | (1,3)=6.115 | 0.0898 |
| AAV-Gq | ARC | intact | 1 | 2 | treatment | (1.524,4.572)=30.04 | **0.0021** |
|  |  |  |  |  | interaction | (2,6)=10.08 | **0.0046** |

GCaMP6f signal was detected in both soma and processes of mCherry-positive cells (Figure 2). The vast majority ofGCaMP6f positive cells were also positive for S100β (94.8±1.2 %, n=3, 5 fields per animal). Calcium transients, characterized by an increase in ΔF/F0, occurred spontaneously in brain slices from all groups. The area under the △F/F0 curves from each individual region of interest (ROI) was used as a measure of the cumulative calcium activity. Bath application of 200nM CNO had no effect in AUC in cells from any mice co-injected with AAV-mCherry control virus regardless of gonadal status or region (Table 3, Figure 2). In gonad intact mice co-injected with AAV-Gq, CNO increased AUC in AAV-Gq infected both the POA and ARC; the increased AUC typically manifested as increased oscillations or oscillations on an elevated plateau as typical of physiologic astrocyte stimulation^43^. In contrast, AAV-Gi had no effect on intracellular calcium in either intact or castrate males.

**Figure 2.**
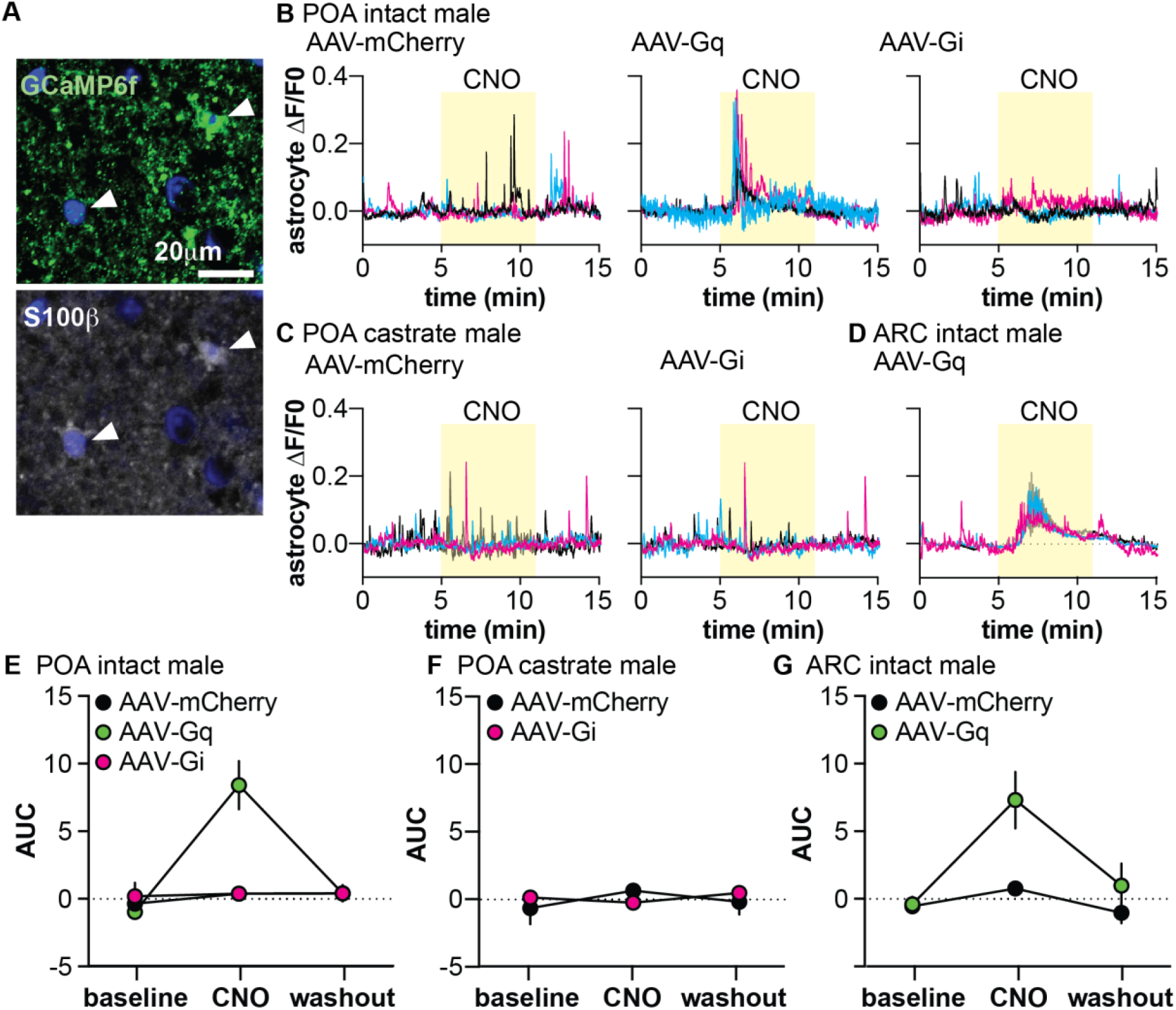
Activation of Gq but not Gi signaling increases intracellular calcium levels in GFAP-expressing cells. A. GCaMP6f (top) and S100β (bottom) are colocalized (arrowheads). B-D. Representative △F/F0 traces from stereotaxic injection of the POA (B,C) or ARC (D). GCaMP6f was co-injected with the virus indicated at the top of each graph. E-G mean ±SEM area under the curve (AUC) for the POA of intact males (E), POA of castrate males (F), and ARC of intact males (G). Two-way repeated-measures ANOVA parameters are in Table 3.

### Gq activation in POA but not ARC GFAP-expressing cells increases LH in vivo

To examine the response of the reproductive neuroendocrine system to activation of Gq signaling in GFAP-expressing cells, serial blood samples obtained from gonad-intact males were assayed for LH. Injection of vehicle had no effect on LH release in any group (Figure 3). When AAV-mCherry control virus was targeted to either the POA or the ARC, CNO injection similarly had no effect on circulating LH levels (Figure 3A, D). When AAV-Gq was targeted to the POA, however, CNO induced an abrupt increase in LH by the next 10-min sample (Figure 3B, C; two-way, repeated-measures ANOVA AAV-mCherry vs AAV-Gq F(1,10)=59, P<0.0001; time F(8,80)=26, P<0.0001; interaction F(8,80)=33, P<0.0001). In marked contrast, neither unilateral nor bilateral hits targeting AAV-Gq to the ARC produced a consistent LH response in the absence of overt neuronal infection (Figure 3E, F; two-way, repeated-measures ANOVA AAV-mCherry vs AAV-Gq F(1,13)=0.1916, P=0.6688; time F(8,104)=1.597, P=0.1345; interaction F(8,104)=0.9265, P=0.4980).

**Figure 3.**
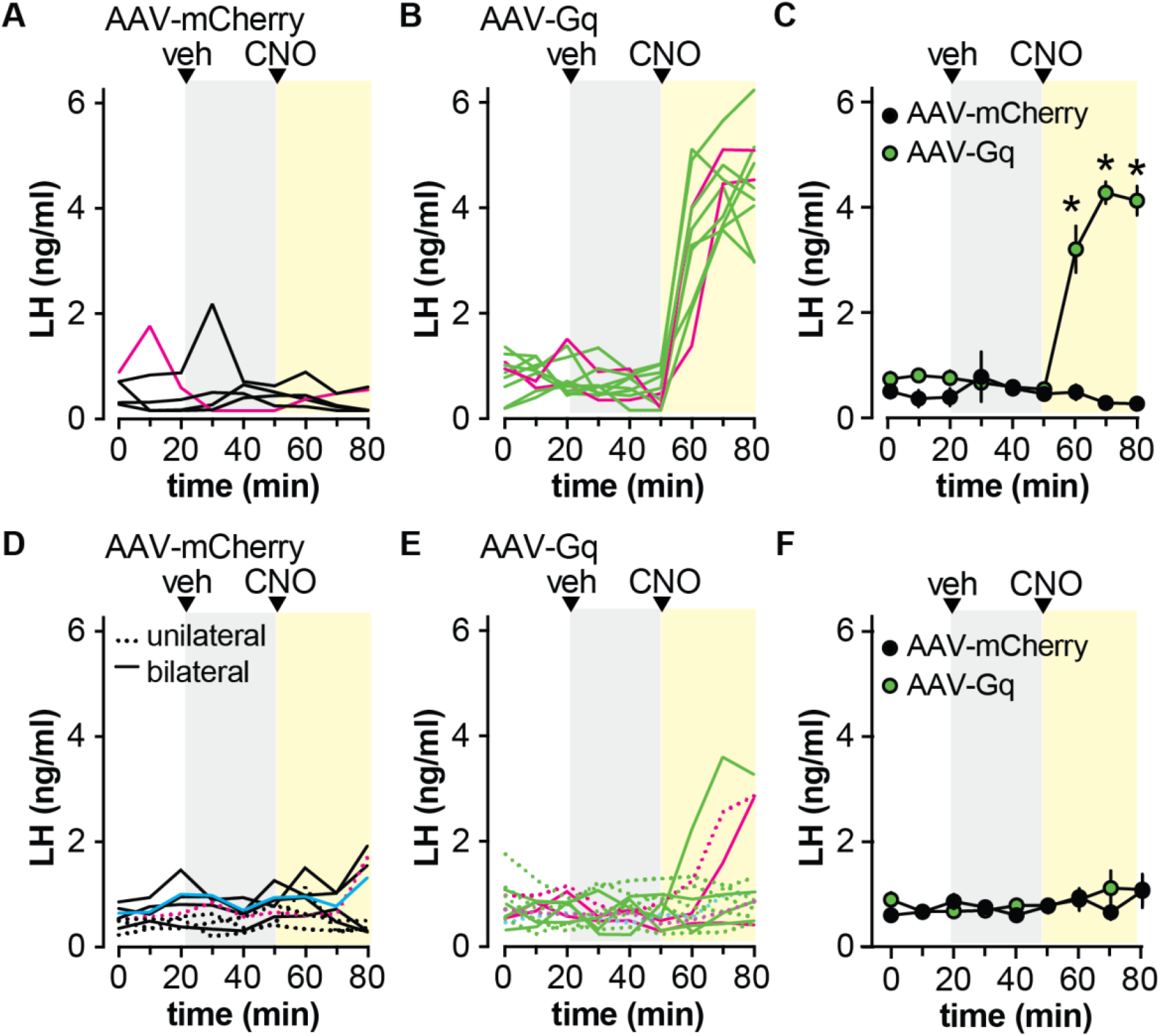
Activation of Gq signaling in GFAP-expressing cells in the POA but not the ARC increases circulating LH. A, B LH levels in individual mice bilaterally injected in the POA with AAV-mCherry (A, black lines) or AAV-Gq (B, green lines). Magenta lines show LH in mice rejected based on infection of unidentified neurons. C. mean ± SEM LH in mice with no observed neuronal infection * P<0.0001 Bonferroni. D, E. LH levels in individual mice with unilateral (dashed lines) or bilateral (solid lines) ARC hits with AAV-mCherry (D, black lines) or AAV-Gq (E, green lines). Magenta lines show LH in mice rejected based on infection of unidentified neurons. F. mean ± SEM LH in mice with no observed neuronal infection. Veh, vehicle

Despite the lack of effect of activating Gi-coupled DREADD with CNO on intracellular calcium levels (Figure 2C,F), we examined LH release in preliminary studies in castrated male mice (n=3/group). CNO had no effect on the mean LH levels (AAV-mCherry, control 4.5±0.7, CNO 3.8±0.5; AAV-Gi, control 4.9±0.3, CNO 4.7±0.5; two-way repeated-measures ANOVA AAV-mCherry vs AAV-Gi F(1,4)=1.4 P=0.2971, control vs CNO (F(1,4)=6.4 P=0.0631), interaction F(1,4)=1.4, P=0.2971). Although the P value for CNO approaches the level accepted for significance, it should be noted this is most likely attributable to values in the AAV-mCherry not the AAV-Gi group. Because these preliminary results were negative for both LH and intracellular calcium with the Gi-coupled DREADD, further experiments were restricted to the Gq-coupled DREADD.

### Gq activation in GFAP-expressing cells of the POA increases GnRH neuron firing rate in vitro

Extracellular recordings of GFP-identified GnRH neurons were made in brain slices from mice infected in the POA with either AAV-mCherry or AAV-Gq. Bath application of CNO (200nM) had no effect on firing rate of GnRH neurons from AAV-mCherry control mice (Figure 4A, D). In contrast, CNO increased firing rate of GnRH neurons within the infected area of AAV-Gq mice as identified by mCherry signal (Figure 4B, D, two-way repeated-measures ANOVA mCherry vs Gq-mCherry F(2, 26)= 4.013, P=0.0303). Interestingly, GnRH neurons located outside of the infected region in AAV-Gq mice, easily distinguishable because they were not surrounded by mCherry signal, did not respond to CNO treatment (Figure 4C). This suggests that activation of Gq signaling in GFAP-expressing POA cells can increase the firing rate of GnRH neurons but that the propagation of that signal to GnRH neurons in uninfected areas is limited.

**Figure 4.**
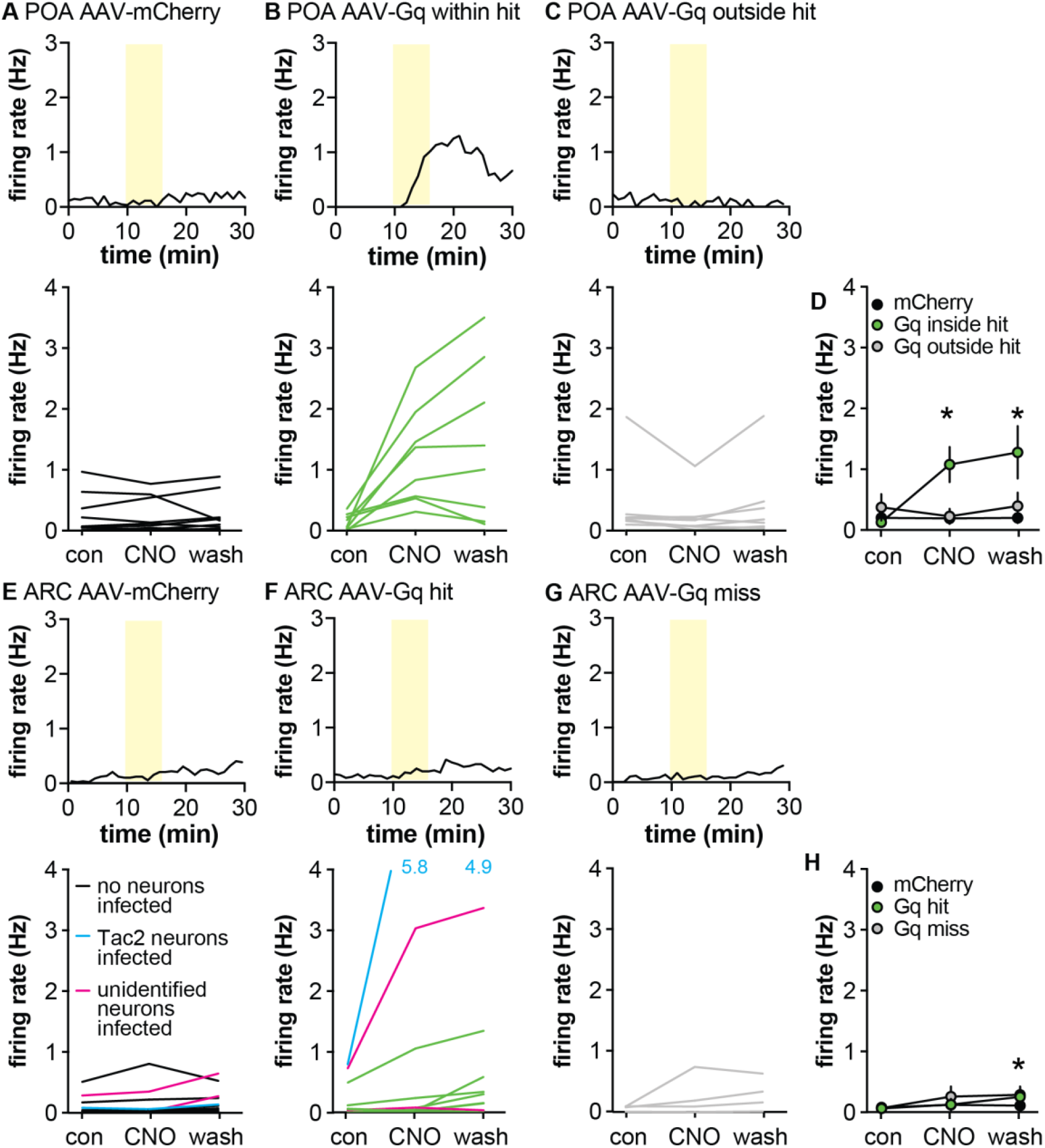
Activating Gq signaling in GFAP-expressing cells differentially affects GnRH vs Tac2 neurons. A-C, top, representative example; bottom firing rate of individual GnRH-GFP neurons in mice injected in the POA with AAV-mCherry (A) or AAV-Gq (B, C). Cells in B were within the mCherry-defined hit, cells in C were outside the hit. D. Mean±SEM firing rate during the different periods. *P<0.0001 Bonferroni Gq inside hit vs Gq outside hit and mCherry. E-G top, representative example; bottom firing rate of individual Tac2-GFP neurons in mice injected in the ARC with AAV-mCherry (E) or AAV-Gq (F, G). Cells in F were within an ARC hit, cells in G were from mice in which the injection missed the ARC. Magenta lines in E and F indicate cells in slices with unidentified infected neurons; cyan lines indicate cells in slices with Tac2 neurons infected. Data from both magenta and cyan cells were excluded from H, mean±SEM firing rate, *P=0.0033 Bonferroni AAV-Gq control vs wash.

To test if activating Gq signaling in GFAP-expressing cells alters firing rate of Tac-2 GFP neurons, the above studies were repeated after targeting injection to the ARC. In this brain region, infection of cells with neuronal morphology within experimental slices was noted. Some of these neurons expressed GFP indicating they are Tac2-expressing neurons, which are known to activate one another ^44^. While these mCherry-expressing neurons were not recorded, they could influence the response within the slice; data from slices with infected neurons were excluded from statistical analyses but are shown in the individual data plots (Figure 4E,F). Neither the type of virus injected (AAV-mCherry vs AAV-Gq) nor the location of the cell inside or outside the hit affected the firing rate of Tac2-GFP neurons in response to bath application of 200nM CNO (Figure 4E-H, Table 4). The firing rate of Tac2-GFP neurons tends to increase over time of recording^45^; this was observed as an increase in firing rate during the wash vs control period.

**Table 4.** Statistical parameters for effects of CNO on neuronal firing rate (Figure 4), P values <0.05 are in bold.

| **GnRH-GFP neurons** | **F** | **P** |  |  |
| --- | --- | --- | --- | --- |
| AAV-mCherry vs AAV-Gq<br>inside hit vs AAV-Gq<br>outside hit | (2,26)=4.013 | **0.0303** |  |  |
| control vs CNO vs wash | (2,52)=7.711 | **0.0012** |  |  |
| Interaction | (4,52)=8.590 | **<0.0001** |  |  |
|  | *Bonferroni* | *con vs CNO* | *con vs wash* | *CNO vs wash* |
|  | AAV-mCherry | >0.9999 | >0.9999 | >0.9999 |
|  | AAV-Gq in hit | **<0.0001** | **<0.0001** | 0.8009 |
|  | AAV-Gq<br>outside hit | >0.9999 | 0>0.9999 | >0.9999 |
| **Tac2-GFP neurons** | **F** | **P** |  |  |
| AAV-mCherry vs AAV-Gq<br>inside hit vs AAV-Gq<br>outside hit | (2,21)=0,2496 | 0.7814 |  |  |
| control vs CNO vs wash | (2,42)=6.751 | **0.0029** |  |  |
| Interaction | (4,42)=1.641 | 0.1819 |  |  |
|  | *Bonferroni* | *con vs CNO* | *con vs wash* | *CNO vs wash* |
|  | AAV-mCherry | >0.9999 | >0.9999 | >0.9999 |
|  | AAV-Gq hit | 0.7680 | **0.0033** | 0.0707 |
|  | AAV-Gq miss | 0.1342 | 0.0626 | >0.9999 |

### CNO-induced increase in GnRH neurons depend at least in part on activation of PGE2 receptors

The putative gliotransmitter PGE2 is primarily produced by astrocytes in the hypothalamus and can increase GnRH neuron firing rate^42,46^ by acting on EP1 and EP2 receptors expressed by these neurons^34^,^47^. We hypothesized that the CNO-induced increase in firing in AAV5-Gq mice is PGE2-dependent. We first examined the effects of the stable PGE2 analogue dimethyl-PGE2 (dmPGE2) on firing rate of GnRH and Tac2 neurons. In GnRH-GFP neurons, pretreatment with either DMSO vehicle (0.3%) or a mix of EP1- and EP2-specific antagonists (100μM SC19220 and 20μM PF04418948, respectively) had no effect on the firing rate (Figure 5A-C). Consistent with previous studies using PGE2^46^, dmPGE2 increased GnRH neuron firing rate in cells pretreated with vehicle for the antagonists (Figure 5A,C, n=9, P<0.0001 two-way repeated-measures ANOVA/Bonferroni, Table 5). Pretreatment with EP1/EP2 receptor antagonists blocked the effect of dmPGE2 on GnRH neuron firing rate (Figure 5B,C, n=10, Table 5). In contrast to GnRH-GFP neurons, neither pretreatment with methylacetate vehicle nor 200nM dmPGE2 had an effect on the firing rate of Tac2-GFP neurons (Figure 5 D-F, n=9 cells, Friedman test F=5.07, P=0.1671).

**Figure 5.**
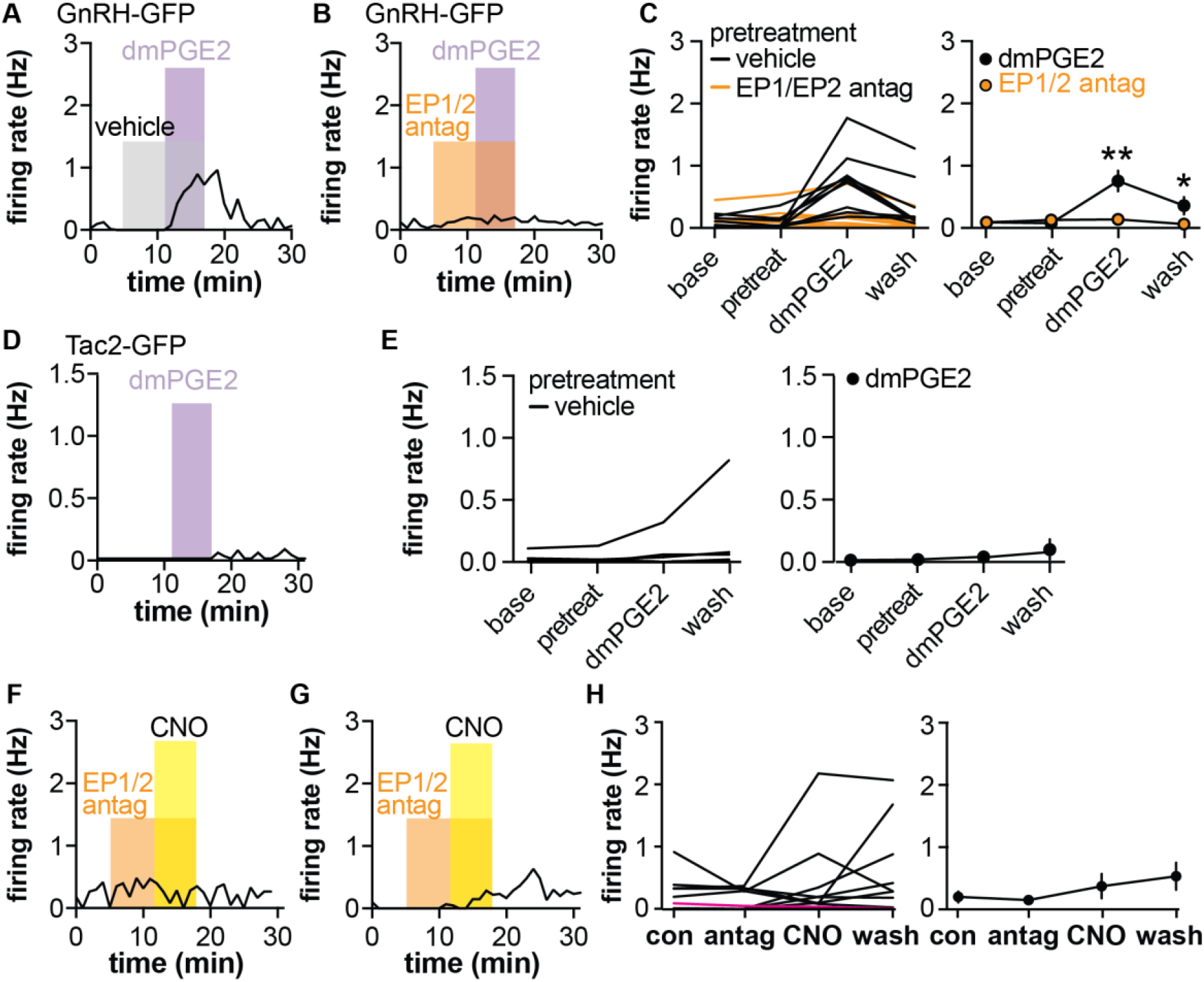
dmPGE2 increases firing rate of GnRH-GFP but not Tac2-GFP neurons; pretreatment (pretreat) with EP1/EP2 receptor antagonists blunts CNO-induced GnRH neuron firing. A, B representative examples of GnRH neuron firing rate response to dmPGE2 following vehicle pretreatment (A) or EP1/EP2 receptor antagonists (B). C, left, mean firing of individual neurons in each period; right group mean±SEM, * P<0.05, ** P<0.0001, two-way repeated-measure ANOVA/Bonferroni. D. representative example of Tac2-GFP neuron response to dmPGE2. Note Y-axis is zoomed in compared to A-C. E, left, Mean firing of individual neurons in each period; right group mean±SEM. F, G Representative examples of GnRH neurons in which pretreatment with EP1/EP2 receptor antagonists blocked (F) or reduced response (G) to CNO. H, left, mean firing of individual neurons in each period, magenta line shows cell in a slice with AAV-Dq infected neurons, which was omitted from the group mean±SEM on the right.

**Table 5.** Statistical parameters for effects of dmPGE2, EP1 and EP2 receptor blockers and CNO on GnRH-GFP neuron firing rate (Figure 5), P values <0.05 are in bold font

| GnRH-GFP neurons | F | P |  |  |
| --- | --- | --- | --- | --- |
| vehicle vs EP1/EP2 antagonist pretreatment | F(1,17) | 0.0229 |  |  |
| time (baseline vs pretreatment vs dmPGE2 vs wash) | F(3,51) | **<0.0001** |  |  |
| Interaction | (3,51)=11 | **<0.0001** |  |  |
|  | *Bonferroni* | *baseline vs pretreatment* | *baseline vs dmPGE2* | *baseline vs wash* |
|  | vehicle | >0.9999 | **<0.0001** | 0.0407 |
|  | EP1/EP2 antagonists | >0.9999 | >0.9999 | >0.9999 |

**Table 6.** Statistical parameters for effects of CNO on GnRH-GFP neuron firing rate in diestrous females (Figure 6), P values <0.05 are in bold font

| GnRH-GFP neurons | F | P |  |  |
| --- | --- | --- | --- | --- |
| AAV-mCherry vs AAV-Gq vs AAV-Gq neurons | (2,23)=45.50 | **<0.0001** |  |  |
| control vs CNO vs wash | (2,46)=45.50 | **<0.0001** |  |  |
| Interaction | (4,46)=30.11 | **<0.0001** |  |  |
|  | *Bonferroni* | *con vs CNO* | *con vs wash* | *CNO vs wash* |
|  | AAV-mCherry | >0.9999 | >0.9999 | >0.9999 |
|  | AAV-Gq | >0.9999 | >0.9999 | >0.9999 |
|  | AAV-Gq with infected neurons | **<0.0001** | **<0.0001** | **0.0023** |

To test if the CNO-induced increase in firing rate of GnRH neurons in brain slices from mice injected with AAV-Dq in the POA was dependent upon PGE2 signaling, slices were pretreated with the EP1/2 antagonist mix before exposure to CNO in the continued presence of the antagonists. Pretreatment with antagonists blunted the CNO-induced increase in GnRH neuron firing (n=11 cells, P=0.4327, Friedman statistic 2.74, Friedman test). This suggests that activating Gq signaling in GFAP-expressing cells stimulates GnRH neurons at least in part via a PGE2-dependent mechanism.

### Activating Gq did not affect reproductive neuroendocrine parameters in vivo or in vitro in females

To examine if the effects of activating Gq signaling in GFAP-expressing cells is sexually-differentiated, females were bilaterally injected in the POA with either AAV-mCherry or AAV-Gq. The survival time post-surgery was longer (3.5-7 vs 18-22 weeks) because of COVID research shut down. LH was measured in 10-min samples taken on diestrus. CNO increased LH in mice infected with AAV-Gq, but all mice exhibited infected cells with neuronal morphology and these data were not considered further (virus F(1,9)=12, P=0.0069; time F(9,81)=21, P,0.0001; interaction F(9,81)=25, P<0.0001). CNO effectively increased intracellular calcium levels in brain slices from females co-injected with AAV-GCaMP6f and AAV-Gq (n=6 slices from 2 mice) but not AAV-mCherry control virus (n=3 slices from 1 mouse; two-way repeated-measures ANOVA virus F(1,7)=4.517 P=0.0712; treatment F(1.497,10.48)=13.44 P=0.0021, interaction F(2,14)=6.550 P=0.0098, Figure 6A).

**Figure 6.**
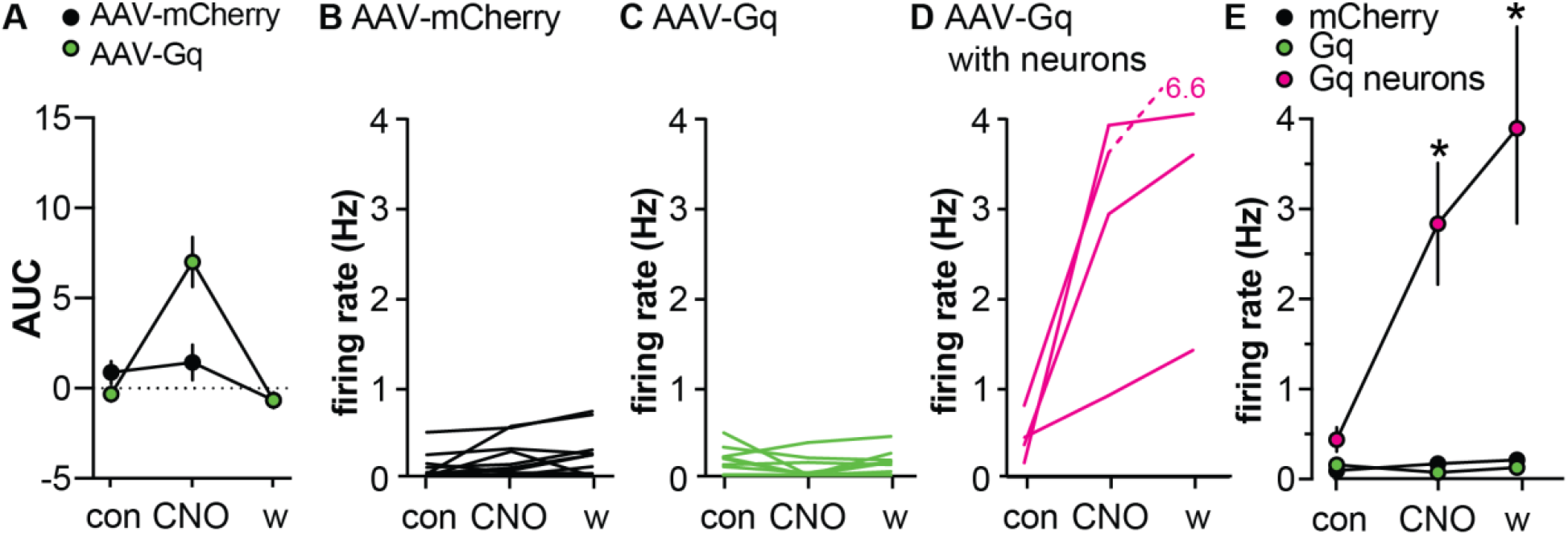
Activating Gq signaling in GFAP-expressing cells in the POA does not affect GnRH neuron firing rate in females. A. Mean ±SEM area under the curve (AUC) of intracellular calcium in cells co-infected with AAV-Gq. B-E Firing rate of individual GnRH-GFP neurons in brain slices from diestrous females injected in the POA with AAV-mCherry (B) or AAV-Gq (C, D); slices in D had viral infection of cells with neuronal morphology. Green symbols and lines show data in AAV-Gq infected slices without detected contamination by neuronal infection; magenta symbols and lines show data in slices with Gq-infected neurons. E. Mean±SEM firing rate during the three periods, *P<0.0001 Bonferroni for cells in slices with Gq-infected neurons vs the other groups.

Despite the ability of CNO to increase intracellular calcium in infected cells, CNO had no effect on firing rate of GnRH neurons in brain slices from female mice infected with either virus (Figure 6B,C,E), unless infected cells with neuronal morphology were observed within the slice (Figure 6D,E). These data suggest there may be a sex difference in the regulation of GnRH neurons by GFAP-expressing cells. An important caveat to point out is that these recordings were made in the afternoon (slices made 2:30-3:30pm, recordings 3:30-9:30pm), whereas recordings from males were made in the morning (slices 9:30am-12pm, recordings 10:30am-6pm). This caveat is mitigated by the consistent virus x treatment interaction for producing elevations in intracellular calcium levels in both sexes. This suggests the lack of GnRH neuron response to CNO in females is mediated subsequent to the GFAP-expressing cell.

## Discussion

The central regulation of fertility depends on the secretion of appropriate patterns of GnRH. An emerging dogma in the field is that this is substantially regulated by arcuate kisspeptin, also known as KNDy, neurons. Here we present evidence that activation of Gq signaling in astrocytes near GnRH neurons in males triggers increased LH release and increased GnRH neuron firing. This is independent of KNDy neurons as similar activation of Gq signaling in astrocytes near these cells fails to alter either LH release or their firing rate. Further, this effect appears to be sexually differentiated as GnRH neurons from diestrous females did not respond. This astrocyte signaling may be a key element in the modulation of GnRH neuron firing and LH release in male mice.

The present findings support and extend previous work suggesting an involvement of astroglia in the regulation of reproductive neuroendocrine function^21,48,49^. Anatomically, a substantial proportion of GnRH neuron somatic membrane is contacted by glia and this contact varies with reproductive state ^22,50^. GnRH terminals are intimately connected with specialized tanycytes in the median eminence; this interaction is dependent upon hormonal milieu in females ^51,52^. Functional interactions among GnRH neurons and glia have been postulated to be primarily mediated by PGE2, as blocking central prostaglandin synthesis reduces gonadotropin release ^53^, whereas injection of PGE2 into the third ventricle or implantation into the preoptic area enhances LH release ^36,54^. More recently, PGE2 was shown to increase firing rate of GnRH neurons in both sexes ^18^. By activating Gq signaling within GFAP-expressing cells directly, the present work extends these findings to the *in vivo* situation in which the entire hypothalamo-pituitary gonadal axis can interact, and indicates a role for these cells within the preoptic area in ultimately increasing LH release in males. In brain slices from these mice, CNO increased intracellular calcium with a similar timecourse to that observed in other brain regions when astrocytes were activated with either DREADDs or by agonists native GPCRs^55,56^. In the present work, the increase in astrocyte intracellular calcium occurred with a shorter lag and had a shorter duration than the changes in neuronal firing rate. This suggests that the GnRH neuron response is dependent upon the elevation of calcium in astrocytes, and also that the signal conveyed to the GnRH neuron produces a prolonged response consistent with activation of a G-protein-coupled receptor within the neurons. This postulate was confirmed as the CNO-induced increase in firing rate response was mimicked by treatment with a stable form of PGE2 and was largely dependent upon activation of EP1 and EP2 receptors.

Astroglia perform several functions within the central nervous system, including release of multiple substances that can serve as gliotransmitters. Astrocytes also are able to communicate over broad areas, propagating elevations in intracellular calcium^29^. This latter observation was key to sparking our interest in how astroglia may function in GnRH release, in particular we postulated that the propagation of signals to distal astrocytes may serve as a mechanism for coordinating activity among the soma of GnRH neurons, which are spread over a wide area from the diagonal band of Broca through the medial basal hypothalamus^57^. It was with this postulate in mind that we recorded from GnRH neurons outside the hit region as defined by mCherry fluorescence. These neurons outside the area of infection did not respond to bath application of CNO with an increase in firing rate, however, suggesting that astroglia do not effectively propagate a signal capable of activating distal GnRH neurons in brain slices.

Interestingly, the activation of Gq signaling in GFAP-expressing cells within the arcuate nucleus was without effect on either LH release or firing rate of KNDy neurons, indicating that the effects on GnRH neurons were independent of this cell type that appears to play a key role in other aspects of GnRH neuron activation. Consistent with a role for PGE2 in the putative astroglia-GnRH communication, treatment of KNDy neurons with PGE2 had no effect upon their firing rate. The lack of effect of astroglia Gq-signaling on KNDy neurons is not attributable to a technical failure as CNO treatment increased intracellular calcium levels in GFAP-expressing cells in this region. Interestingly, the area infected by virus includes tissue through which the processes of GnRH neurons pass *en route* to the median eminence. This suggests that this region of the GnRH neuron axon may not be sensitive to mediators released by astroglia. Of interest in this regard, treatment of explants containing the median eminence along with the medial basal hypothalamic region with PGE2 induced GnRH release ^58^. Our arcuate hits typically did not extend into the median eminence itself. As was the case with GnRH neurons outside the hit, it is possible there is not sufficient propagation of astroglial signals effective in activating GnRH neuron terminals outside the local region of infection. It is also possible that astrocytes exhibit regional variation in the gliotransmitters they produce. Of note, however, activation of Gq signaling in these cells would be postulated to induce release of whatever substances are downstream of that signaling pathway in the cells and not be limited to prostaglandin synthesis. Together these observations suggest the effects of Gq signaling within astroglia are at least in part specialized to preoptic regions of GnRH neurons for the reproductive neuroendocrine system.

We conducted limited studies of activating Gq signaling in astroglia in diestrous females. CNO treatment effectively induced increases in intracellular calcium, but failed to alter GnRH neuron firing rate unless there were overtly infected neurons within the slice. There are several possible explanations for these observations. First cycle stage may affect the response, although the elevation of intracellular calcium in the present study suggests that astroglia were activated in the diestrous mice studied. Second different gliotransmitters may be released upon astroglia activation in females, perhaps also dependent upon cycle stage. In prior work, over two-thirds of GnRH neurons increased firing rate in response to PGE2 regardless of sex or of cycle stage in females ^18^, indicating responsiveness to this mediator is consistent among groups. Third, it is possible that the relevant astroglia regulating GnRH release are located in different brain regions, for example more caudal within the anteroventral periventricular region known to be important for induction of the female-specific preovulatory GnRH and LH surge^59^.

In contrast to the robust serum LH and GnRH neuron firing response to activation of Gq signaling in astroglia, activation of Gi signaling had no effect under the conditions studied. This included no effect on intracellular calcium levels in POA astroglia in either intact or castrated mice, and no effect on LH in castrated mice. The effects of activating Gi signaling in GFAP-expressing cells varies with brain region. In the hippocampus, increases in intracellular calcium were reported^60^, whereas in the hypothalamic arcuate region, activation of Gq and Gi signaling have opposing effects on food intake^41^. As in females, interpretation of lack of effect must be limited to the experiments conducted, and it is possible that activation of Gi signaling in astroglia in other brain regions will alter the output of GnRH and LH. Interestingly, there are no reports of receptors acting via Gi-coupled receptors within the preoptic area/hypothalamus.

In the present work, the main mediator of the effects of activating Gq signaling in astroglia appeared to be PGE2. GnRH neurons express both EP1 and EP2 receptors^34^, thus PGE2 could be acting directly upon these cells. There are several things we cannot rule out from our studies. First, while astroglia appear to be the primary source of PGE2 in the hypothalamus, it is possible that other cell types also produce this signal. Second, it is possible that mediators other than PGE2 are produced by astroglia in response to activating Gq signaling. In this regard, in addition to prostaglandins, these cells have been shown to release other mediators including glutamate^61–63^ and ATP ^64,65^. Finally, any of these mediators may act via intermediate cell types within the brain slice to bring about effects on GnRH neuron firing. An intriguing question that remains to be answered is what substances serve as the endogenous activators of Gq signaling in POA astroglia. Activation of group 1 metabotropic glutamate receptors induces release of transforming growth factor alpha and neuregulins, which activate ErbB receptors on astroglia. This activation leads to activation of cyclooxygenase 2 (COX-2) a rate limiting step in the synthesis of PGE2 from arachidonic acid. While pure speculation, there are a number of interesting substances active within the reproductive neuroendocrine system that act via Gq-coupled receptors, including neurokinin B, kisspeptin and GnRH itself.

In summary, the present work defines that activation of Gq signaling in astrocytes can increase GnRH neuron activity and LH release in a sex-dependent manner. Activation of this pathway may activate this system independent of arcuate kisspeptin neurons or may sculpt the response to that or other critical inputs to GnRH neurons to affect ultimately reproduction.

## Materials and Methods

All reagents were purchased from Sigma-Aldrich (St. Louis, MO, USA) unless noted.

### Animals

Mouse strains used for this work are summarized in Table 1. Mice expressing enhanced green fluorescent protein (GFP) under the control of GnRH promoter (GnRH-GFP mice, JAX 033639)^66^ or Tac2-GFP BAC transgenic mice (015495-UCD/STOCK Tg [Tac2-EGFP]381Gsat, Mouse Mutant Regional Resource Center (http://www.mmrrc.org/))^67^ were used to identify GnRH and KNDy neurons for recording, respectively. *Tac2* encodes neurokinin B, which is co-expressed with kisspeptin and dynorphin in KNDy neurons. Tac2-GFP-identified cells in brain slices used for recording also express kisspeptin and/or dynorphin at high percentages, supporting their identity as KNDy neurons^67^. Mice were held on a 14h light/10h dark light cycle with lights on at 0300 Eastern Standard Time and had *ad libitum* access to water and chow (Teklad 2916). Adult gonad-intact males and females were used between 60 and 220 days of age. In females, estrous cycle stage was determined by vaginal lavage and confirmed by uterine mass (≥80mg for diestrus)^68^. Castration efficacy in males was determined by seminal vesicle mass (intact >250mg, castrate <150mg). The Institutional Animal Care and Use Committee at the University of Michigan approved all procedures.

### Stereotaxic injections

Viruses used in this work are summarized in Table 1. Mice were anesthetized with isoflurane to effect and received 5mg/kg carprofen before the surgery and 24h post-surgery for analgesia. AAV5 carrying a payload encoding the hM3Dq (Addgene 50478) or the hM4Di (Addgene 50479) DREADDs or control AAV (Addgene58909) fused to mCherry under the control of the GFAP-promoter were used to introduce DREADDs to regions of interest. AAV5 encoding GCaMP6f under the control of GFAP promoter (Addgene 52925) was used to target this calcium indicator to GFAP-expressing cells. AAVs (50-100nL) were administered by bilateral stereotaxic injection into the POA (AP: −2.27mm; ML: −0.3 and +0.3mm; DV: −4.55mm from frontal vein) or ARC (AP: −1.6mm; ML: −0.2 and +0.2mm; DV: −5.95mm from Bregma). Mice were monitored until fully recovered from anesthesia and surgery sites were examined daily for 10d. Viral infection was allowed to proceed for 3-6wks before experiments.

### Immunohistofluorescence

Mice were deeply anesthetized with isoflurane and transcardially perfused with 0.9% NaCl (10mL) then 10% neutral-buffered formalin for 15min (~50mL). Brains were placed into 10% formalin for 4h and transferred into 20% sucrose in 0.1M PBS for cryoprotection for at least 48h and until sectioning. Four series of 30μm free-floating sections were obtained with a cryostat (Leica CM3050S) in 0.1M phosphate-buffered saline (PBS) pH=7.4, then transferred io antifreeze solution (30% ethylene glycol, 20% glycerol in PBS) for storage at −20°C. Sections were washed three times in PBS, incubated in blocking solution (PBS containing 0.4% Triton X-100, 2% normal goat serum, Jackson Immunoresearch) for 1hr at room temperature, and incubated in primary antibody (Table 2) diluted in blocking solution for 48h at 4°C. Sections were washed three times in PBS and incubated with Alexa-conjugated secondary antibodies for 1.5h at room temperature (Molecular Probes and Jackson Immunoresearch, 1:500). After 3 washes with PBS, slices were incubated with 300nM 4’,6-diaminidino-2-phenylindole dihydrochloride (DAPI) in PBS for 10min at room temperature. Slices were washed 3 times in PBS, mounted on Superfrost plus slides (Fisher Scientific) with ProLong Gold antifade reagent (Invitrogen) and coverslipped (VWR International). Primary antibodies and dilutions used are in Table 2. Images were collected on a Zeiss AXIO Imager M2 (lower magnification) or on a Nikon A1 confocal microscope (colocalization studies). The number of mCherry only, mCherry/S100β, and mCherry/NeuN coexpressing cells was counted from confocal pictures (3.49μm optical sectioning, same exposure for each signal, levels adjusted to represent the signal observed by eye, five fields/mouse in the infected region; AAV-mCherry and AAV-Gq n=3 mice each POA and ARC, AAV-Gi n=4 mice POA). Brightness was increased 30% in pictures used for figures. In addition to immuno-identification, all 300μm slices for imaging and electrophysiology were examined for infection of cells with neuronal morphology. Data from slices or mice in which infection of neuronal cells was detected were eliminated from statistical analysis but individual data are shown for transparency in the figures.

### Tail tip blood collection

To examine the effect of DREADD activation on LH levels, mice were handled daily for 2wks before CNO administration studies or 5wks before sampling LH pulses and habituated to IP injection of 0.9% saline for the last 3-4d. The tip of the tail was nicked and 6μL of blood was collected and mixed immediately with 54μl of assay buffer (PBS, 0.05%Tween and 0.2% BSA). Sampling regimen and animal models were selected based on postulated LH response to activating Gq or Gi signaling within GFAP-expressing cells. To test the effects of activating Gq, postulated to be activating, gonad-intact males were sampled every 10min for 2h, with IP injection of 0.9% saline vehicle at 30min, and IP injection of 0.3mg/kg CNO at 60min. To test the effect of activating Gi, postulated to be inhibitory, males were castrated to elevate LH release; one week after castration, these mice were sampled every 6min for 3h, with IP injection of 0.3mg/kg CNO at 90min.

### LH assay

Tail blood diluted with assay buffer was kept on ice until the end of sampling then stored at −20°C until LH assay by the University of Virginia Ligand Assay and Analysis Core^48^. The capture monoclonal antibody (anti-bovine LHβ subunit, 518B7) is provided by Janet Roser, University of California, Davis. The detection polyclonal antibody (rabbit LH antiserum, AFP240580Rb) is provided by the National Hormone and Peptide Program (NHPP). HRP-conjugated polyclonal antibody (goat anti-rabbit) is purchased from DakoCytomation (Glostrup, Denmark; D048701-2). Mouse LH reference prep (AFP5306A; NHPP) is used as the assay standard. The limit of quantitation (functional sensitivity) was 0.016 ng/ml, defined as the lowest concentration that demonstrates accuracy within 20% of expected values. Coefficient of variation (%CV) was determined from serial dilutions of a defined sample pool. Intraassay CV was 2.2%; interassay CVs were 7.3% (low QC, 0.13 ng/mL), 5.0% (medium QC, 0.8 ng/mL) and 6.5% (high QC, 2.3 ng/mL).

### Brain Slice Preparation

All solutions were bubbled with 95% O2/5% CO2 for at least 30 min before exposure to tissue and throughout the experiments. At least one week after sampling for LH, brain slices were prepared through the hypothalamus as described^69^. The brain was rapidly removed and placed in ice-cold sucrose saline solution containing (in mM): 250 sucrose, 3.5 KCl, 26 NaHCO3, 10 glucose, 1.25 Na_2_HPO_4_, 1.2 MgSO_4_, and 3.8 MgCl_2_. Coronal slices (300μm) were cut with Leica VT1200S (Leica Biosystems, Buffalo Grove, IL, USA). Slices were incubated for 30min at room temperature (~21-23 °C) in 50% sucrose saline and 50% artificial cerebrospinal fluid (ACSF) containing (in mM) 135 NaCl, 3.5 KCl, 26 NaHCO3, 10 D-glucose, 1.25 Na_2_HPO_4_, 1.2 MgSO_4_, 2.5 CaCl_2_ (pH 7.4), then transferred to 100% ACSF at room temperature for 0.5-4.5h before recording. At the end of the recording, slices of interest were fixed with 10% formalin for 40min, washed three times in PBS, and were processed for immunofluorescence to study the spread of the infected area or examined directly for mCherry signal in cells with neuronal morphology.

### Electrophysiological recordings

Targeted single-unit extracellular recordings were used to minimize impact on the cell’s intrinsic properties^70^’^71^. Recording pipettes (2-4MΩ) were pulled from borosilicate glass (Schott no. 8250; World Precision Instruments, Sarasota, FL, USA) with a Sutter P-97 puller (Sutter Instrument, Novato, CA, USA). Pipettes were filled with HEPES-buffered solution containing (in mM): 150 NaCl, 10 HEPES, 10 glucose, 2.5 CaCl_2_, 1.3 MgCl_2_, and 3.5 KCl, and low-resistance (6-15MΩ) seals were formed between the pipette and neuron. Recordings were made in voltage clamp with a 0mV pipette holding potential, data acquired at 10kHz and filtered at 5kHz using a one amplifier of an EPC10 dual patch clamp amplifier controlled with PatchMaster software (HEKA Elektronic, Lambrecht, Germany).

Slices were transferred to a recording chamber with constant perfusion of carboxygenated ACSF at 29-32°C at a rate of approximately 3ml/min. GFP-positive cells were targeted for recording in the POA (GnRH neurons) or ARC (KNDy neurons). Cells that were surrounded by mCherry signal were considered within the infected area (hit) and those with no detectable peripheral mCherry signal within 150μm were considered outside the infected area (miss). Recordings consisted of a 5-10min stabilization period, a 5-min control period, bath-application of treatment for six minutes, followed by a wash. Mean firing rate was calculated for the last 3-min baseline period, for min 5-7 of treatments, and min 8-10 after treatment (wash). At the end of each recording, cells that were inactive throughout the recording were treated with 20 mM potassium in ACSF; cells that exhibited action currents in response were verified to be alive and recordable, and all data were used. Cells that did not respond to elevated potassium were excluded. No more than three cells from the same animal were included and at least four animals were studied per group for electrophysiology studies.

To test the hypothesis PGE2 affects GnRH and KNDy neuron firing, we used 16,16-dimethyl prostaglandin E2 (dmPGE2, 200nM), a stable form of PGE2 (Cayman Chemical 14750) in 0.00076% methyl acetate vehicle. Antagonists for prostaglandin receptor EP1 (SC-19220, 100μM) and EP2 (PF-04418948, 20μM) were diluted into fresh bubbled ACSF (final DMSO concentration 0.3%) and bath applied before CNO treatment. The recording paradigm was a 6min baseline, 6-min pretreatment with methyl acetate vehicle containing either DMSO vehicle or EP1 and EP2 antagonists, a 6-min treatment that added 200nM dmPGE2, followed by wash in ACSF. Mean firing rate was calculated for the last 3-min baseline and pretreatment periods, minutes 5-7 following addition of dmPGE2 for treatment, and min 8-10 after wash began.

### Calcium imaging

Calcium imaging was performed on brain slices from mice co-injected with AAV5-gfaABC1D-cyto-GCaMP6f and either AAV-mCherry control, the AAV-Gq or the AAV-Gi (ratio 1:1). Calcium was imaged for a 50ms exposure time at 0.5s intervals for 20min with an Hammamatsu ORCA-FLASH4.0 V3 CMOS camera. Imaging began after a 5-min stabilization period (not included in the analysis), and consisted of a 5-min control period, a 6-min treatment with 200nM CNO, and 9-min wash. Regions of interest (ROIs) were isolated from astrocytic domains for which signal fluctuations were identified prior to CNO treatment. Each ROI likely represents the soma or proximal regions of a single astrocyte. For each slice, a background ROI was isolated from a region in which no signal fluctuations were detected; a single region was used as the background reference for all the ROIs of a slice. ΔF/F0 [(F(t) −F0(t)) / F0(t)] was calculated using custom routines in Igor Pro (Wavemetrics) and plotted as a function of time for each ROI. Briefly, the background ROI was subtracted from all other ROIs from that slice. To account for the signal decay related to photobleaching, we used a double exponential curve fit for each individual ROI that was considered F0; the CNO treatment period was excluded from the curve fit using the masking capability in the Igor Pro curve fitting function. We used the resulting curve fit as an estimate of F0 as a function of time. Each curve fit was verified by eye to conform to the timecourse of the bleaching for each ROI. F0 was subtracted from the background-subtracted traces to calculate △F. △F was divided by the fit to obtain ΔF/F0. To quantify the effect of CNO on the astrocytic calcium signals, the area under the curve (AUC) from each ΔF/F0 curve was calculated for different periods (control, the last 3-min baseline period; CNO from the first 3min of CNO exposure; wash AUC from min 5-7 after CNO treatment ended. The AUCs obtained from 27 independent ROIs were average to obtain the average AUCs of the slice (n=1). Three to six slices from 1-2 mice were used for each group (Table 3).

### Analysis and statistics

Data were analyzed using code written in IgorPro 8 (Wavemetrics). Action currents (events) were detected and confirmed by eye. Statistical analyses were conducted with Graphpad Prism 9. Distribution normality was checked with Shapiro-Wilk; most experiments included one or more groups that were not normally distributed. One-way repeated-measures tests were conducted with the Friedman test. For two-way repeated-measures ANOVA the Bonferroni *post hoc* is considered sufficiently rigorous for non-normally distributed data^72^. Statistical tests for each study are specified in the Results. P<0.05 was considered significant and exact P-values are reported.

## Acknowledgements

We thank Elizabeth Wagenmaker and Laura Burger for expert technical assistance.

## Abbreviations

GnRH: gonadotropin-releasing hormone;
LH: luteinizing hormone;
KNDy: neurons co-expressing kisspeptin, neurokinin B and dynorphin;
POA: preoptic area;
ME: median eminence;
ARC: hypothalamic arcuate nucleus;
GFAP: glial fibrillary acidic protein;
(AAV): adenoassociated virus;
GFP: green fluorescent protein;
CNO: clozapine n-oxide;
PGE2: prostaglandin E2;
dmPGE2: dimethyl prostaglandin E2;
DREADD: designer receptor exclusively activated by designer drug;
GPCR: G-protein-coupled receptor

---

**Supplementary Figure S1.**
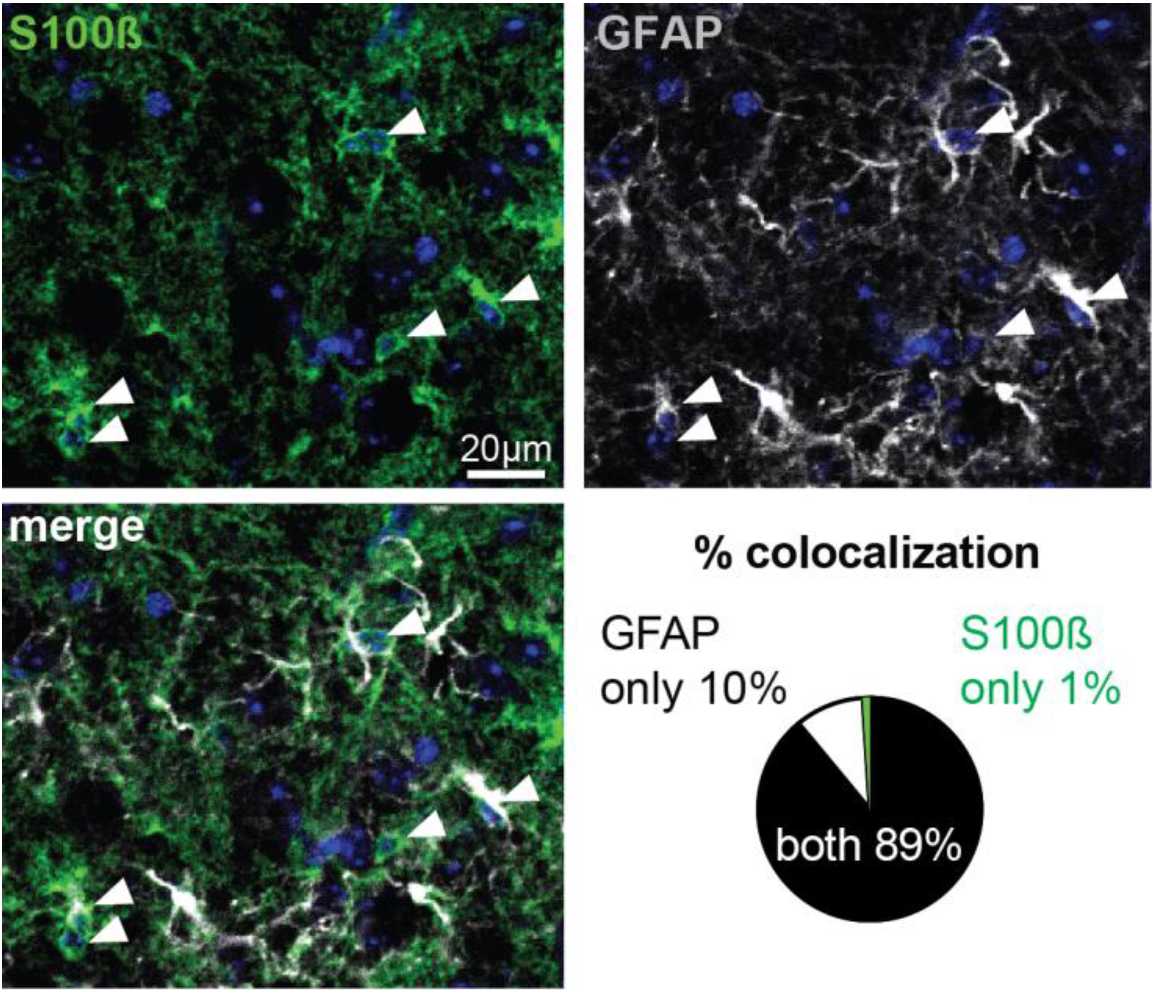
GFAP and S100β signals are co-expressed in the majority of cells (n=3 mice, 5 fields/mouse).

